# Targeting neuronal homeostasis to prevent seizures

**DOI:** 10.1101/2022.03.07.483229

**Authors:** Fred Mulroe, Wei-Hsiang Lin, Connie Mackenzie-Gray Scott, Najat Aourz, Yuen Ngan Fan, Graham Coutts, R Ryley Parrish, Ilse Smolders, Andrew Trevelyan, Robert Wykes, Stuart Allan, Sally Freeman, Richard A. Baines

## Abstract

Manipulating neuronal homeostasis, which enables neurons to regulate their intrinsic excitability, offers an attractive opportunity to prevent seizures. However, no anticonvulsant compounds have yet been reported that directly manipulate neuronal homeostasis. Here, we describe a novel class of anticonvulsant compounds, based on 4-*tert*-butyl-benzaldehyde (4-TBB), with a mode-of-action that includes increased expression of the homeostatic regulator Pumilio (Pum). In *Drosophila* and mouse we use a pentylenetetrazole (PTZ) induced seizure model, and an electrically induced seizure model for refractory seizures to evaluate anticonvulsant efficacy. The pyrazole analogue (RAB216) demonstrates best efficacy, protecting 50% of mice from PTZ-induced seizure. Knock-down of Pum, in *Drosophila*, blocks anticonvulsive effects, whilst analysis of validated Pum targets show significant reductions following exposure of mouse brain to 4-TBB. This study provides proof-of-principle that anticonvulsant effects can be achieved through regulation of neuronal homeostasis and identifies a chemical lead compound for future development.

## Introduction

Neuronal homeostasis provides an attractive target to achieve therapeutic control of epilepsy. This is because homeostatic mechanisms ostensibly oppose extremes of neuronal activity that are associated with seizures. By maintaining neuronal activity patterns at physiologically relevant ‘set-points’, neuronal homeostasis acts to ensure stability of both neuron and network function across the life-course ^1^. However, to date, neuronal homeostasis has not been specifically targeted for clinical benefit.

Pumilio (Pum) homeostatically maintains action potential firing rates within a set-range ^2^. A translational repressor, Pum binds mRNA transcripts and reduces *de novo* protein synthesis, with increased Pum expression occurring in neurons exposed to increased synaptic excitation. Conversely, as synaptic excitations fall, Pum expression is reduced ^3^. The 3’-UTR of Pum-regulated transcripts usually contain one or more copies of a Pum-Response Element (PRE: UGUANAUA, where N is A, C, G or U) ^4^. Analysis of both *Drosophila* and mammalian transcriptomes identifies more than 1000 transcripts that contain one or more PREs, consistent with a broad regulatory role ^5^. However, Pum requires additional co-factors (including Nanos and Brain-tumor) and the actual effect of Pum is likely dictated by both the number and proximity of these additional binding elements, in addition to the number of PREs ^6,7^. In mammals, regulated transcripts that have potential to influence neuron activity include Na_v_1.6 (SCN8A) ^2^ and GLUR2 (AMPA receptor) ^8^. Pum-dependent homeostatic translational repression of Na_v_1.6, in rat cortical pyramidal neurons, reduces the amplitude of expressed voltage-gated Na^+^ current (I_Na_) and lowers action potential firing frequency ^2^. Down-regulation of AMPA receptor expression may also be expected to be antiepileptic, evidenced by the anti-epileptic compound, perampanel, which is an allosteric antagonist of AMPA receptors ^9^.

Whilst *Drosophila* has one *pum* gene, mammals express two highly similar variants (*pum 1 & 2*) that are co-expressed, and which bind identical RNA motifs and, thus, appear to act redundantly ^10-12^. Seizure occurrence could reflect reduced homeostatic capability and it is significant that recent studies suggest reduced Pum contributes to epilepsy. Specifically, i) *pum1* or *2* haploinsufficiency is associated with spontaneous seizures in mice ^13-15^, ii) Pum2 expression is reduced in human patients suffering temporal lobe epilepsy and in rat hippocampus following pilocarpine-induced seizure ^16^, and iii) *dpum* expression is reduced in *Drosophila* genetic seizure mutations ^17^. In the latter, transgenic up-regulation of dPum is potently anticonvulsant in these same *Drosophila* mutations ^17^. Based on a screen to identify chemicals that increase expression and/or stability of dPum, we identified avobenzone, which secondary screens show is anticonvulsant in seizure-sensitive *Drosophila* ^17^. However, the physiochemical properties of avobenzone are not compatible with clinical use. Thus, in this study, we report the identification of an avobenzone analogue, 4-*tert*-butyl-benzaldehyde (4-TBB), that is anticonvulsant and has properties more consistent with clinically active compounds. We show that 4-TBB and analogues (specifically the pyrazole RAB216) are active against a range of *Drosophila* seizure mutants and, significantly, reduce severity of both pentylenetetrazol (PTZ)-and pharmacoresistant electrically (6Hz)-induced seizures in mouse. Reduction of seizures, in fly and mouse, is accompanied by increased expression of Pum. We further report down-regulation of known Pum targets following exposure of mouse brain to 4-TBB thus validating proposed mode-of-action.

## Results

### 4-TBB suppresses seizure behaviour in *Drosophila*

Single *Drosophila* gene mutations increase seizure-like activity in response to electrical shock ^18^. In a prior study, we identified avobenzone to both increase *dpum* expression and reduce seizure severity in *Drosophila* ^17^. Avobenzone is poorly soluble and therefore we identified a breakdown product, 4-TBB to be a better candidate for analysis of mode-of-action. We tested the anticonvulsant activity of 4-TBB in three diverse seizure mutants to demonstrate wide applicability. Exposure to drug (1.2 mM contained within food) was sufficient to reduce electroshock-induced seizure duration in *bangsenseless*^*1*^ (*para*^*bss*^; Na_v_ hypermorph, 194 ± 116 vs. 348 ± 101 s, 4-TBB vs. control, n = 30, *p* < 0.001), *easilyshocked* (*eas*; ethanolamine kinase deficiency, 160 ± 81 vs. 232 ± 124 s, n = 30, *p* = 0.011) and *julius seizure* (*jus*; a transmembrane domain protein of undetermined function, 197 ± 124 vs. 266 ± 112 s, n = 30, *p* = 0.03, Fig. 1A). The level of seizure suppression observed for 4-TBB, in all mutations, is favourable compared to phenytoin (PHE, *c*.*f*. Fig. 1D).

**Figure 1.**
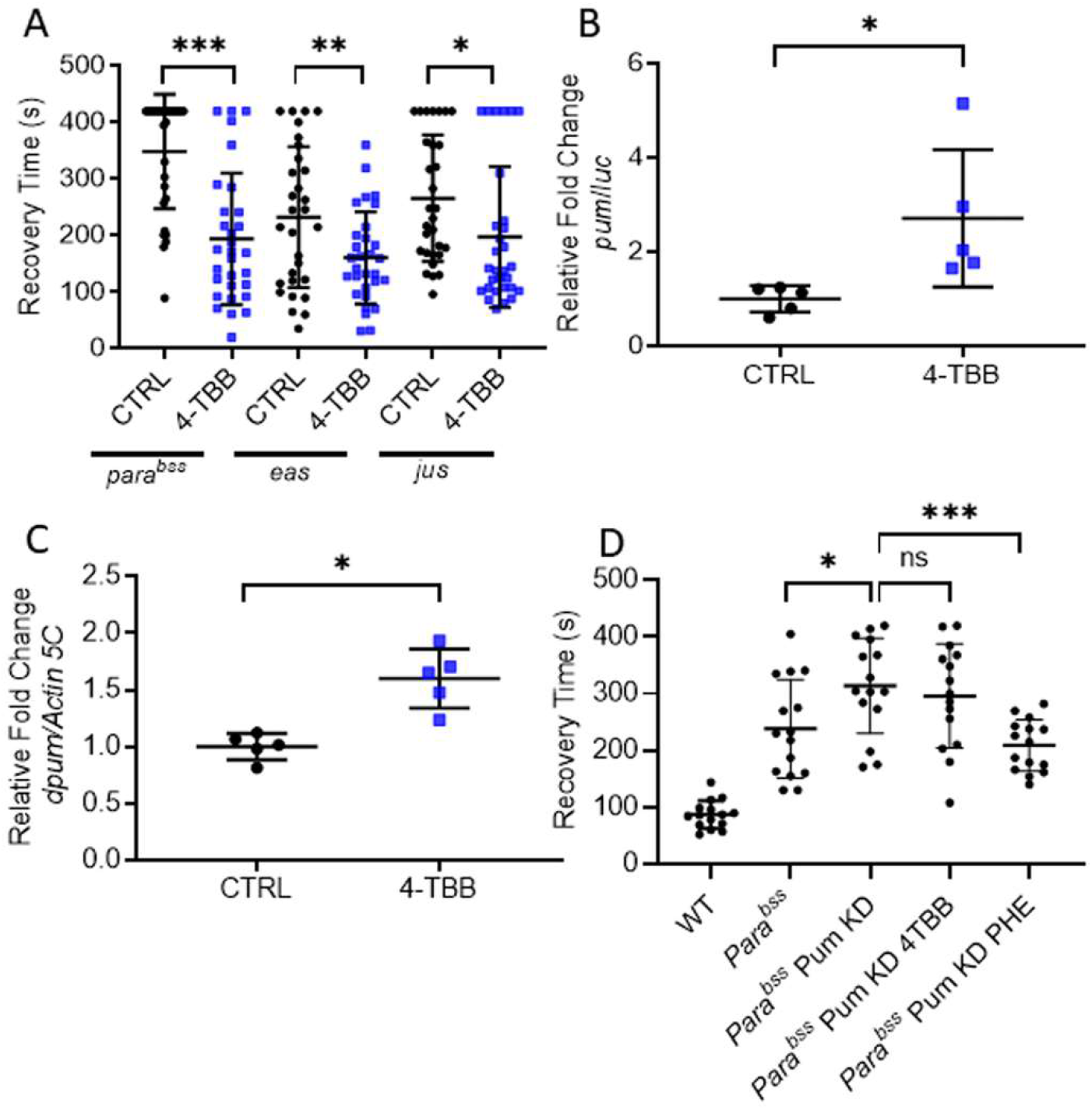
4-TBB is anticonvulsant in *Drosophila*. **A**) An anticonvulsant effect of 4-TBB is present in three independent *Drosophila* seizure mutations, *para*^*bss*^ (*p* < 0.001), *eas* (*p* = 0.011) and *jus* (*p* = 0.03) (unpaired *t*-tests, n = 30 larvae for each treatment). **B**) Identical exposure of transgenic *dpum* promoter-GAL4>UAS-Luciferase (Luc) larvae to 4-TBB results in significant up-regulation of Luc expression (*p* = 0.03, unpaired *t*-test, n = 5 independent samples / 10 CNS per sample). **C**) Ingestion of 4-TBB is sufficient to increase *dpum* mRNA abundance measured by QRT-PCR (*p* = 0.002, unpaired *t*-test, n = 5 independent samples / 20 CNS per sample). **D**) RNAi-mediated knockdown of *dpum* expression in *para*^*bss*^ expectedly increases seizure recovery time ^17^ (*p* = 0.03). Exposure to 4-TBB is ineffective in this background (*p* = 0.87), whilst exposure to phenytoin (PHE, 2mM in food) remains anticonvulsant (*p* = 0.002). Significance tested using a one-way ANOVA (F_(4, 70)_ = 24, *p* = 0.0009) followed by Dunnett’s post-hoc test (multiple experimental groups). Wildtype (WT) recovery time is shown for comparison. Bars report means ± SD (n = 15). * = *p* < 0.05, ** = *p* < 0 .01, *** = *p* < 0.001, ns = not significant.

The expression of *dpum* is reduced in *para*^*bss*^ mutants and, moreover, increasing *dpum* expression in this mutant background is sufficient to suppress seizure activity in response to electroshock ^17^. To determine whether the anticonvulsant effect of 4-TBB is associated with up-regulation of *dpum*, we used a *dpum*-minimal promoter construct upstream of GAL4 (*dpum*-GAL4) to drive expression of UAS-luciferase (UAS-luc) ^19^. Exposure of *dpum*-GAL4>UAS-Luc flies to 4-TBB (1.2 mM in food) resulted in a significant increase in luciferase activity (2.7 ± 1.5 fold change, n = 5, *p* = 0.03, vehicle control set as 1) (Fig. 1B). We adopted this approach because available anti-Pum antibodies (designed to rodent Pum1 and Pum2) do not work well in *Drosophila*. We also observed a significant increase in *dpum* transcript abundance, measured by QRT-PCR, of ∼60% (1.6 ± 0.3 fold change, n = 5, *p* = 0.002, vehicle control set as 1) following exposure to 4-TBB (Fig. 1C). Finally, we find that the anticonvulsive activity of 4-TBB is significantly diminished when *dpum* expression is reduced in the CNS, via targeted expression of an RNAi construct (Fig. 1D). PHE, which has a different mode of action ^20^, remains active under these conditions. We conclude that the mode-of-action of 4-TBB requires the presence of dPum and increases expression of this homeostatic regulator.

### 4-TBB reduces epileptiform activity in mouse hippocampal culture slices

Incubation of acutely-harvested mouse brain slice with 4-TBB (1.2mM) for 2h was sufficient to produce a significant increase in Pum2 expression (Pum1 not measured), as determined by Western Blot (2.5 ± 0.3 fold compared to vehicle-treated controls which were the corresponding slice taken from the opposite hemisphere incubated in vehicle only, set at 1, n = 5, *p* = 0.002, data not shown). Addition of 4-TBB (1.2mM) to mouse organotypic hippocampal slices, in which epileptiform activity was already established (Fig. 2A-B), led to a progressive reduction in epileptiform activity (*p* < 0.0001; n = 7 and 8, control vs. 4-TBB, respectively) compared to controls (vehicle added) that was apparent by 1h, and maximal after 3h (Fig. 2C-D).

**Figure 2.**
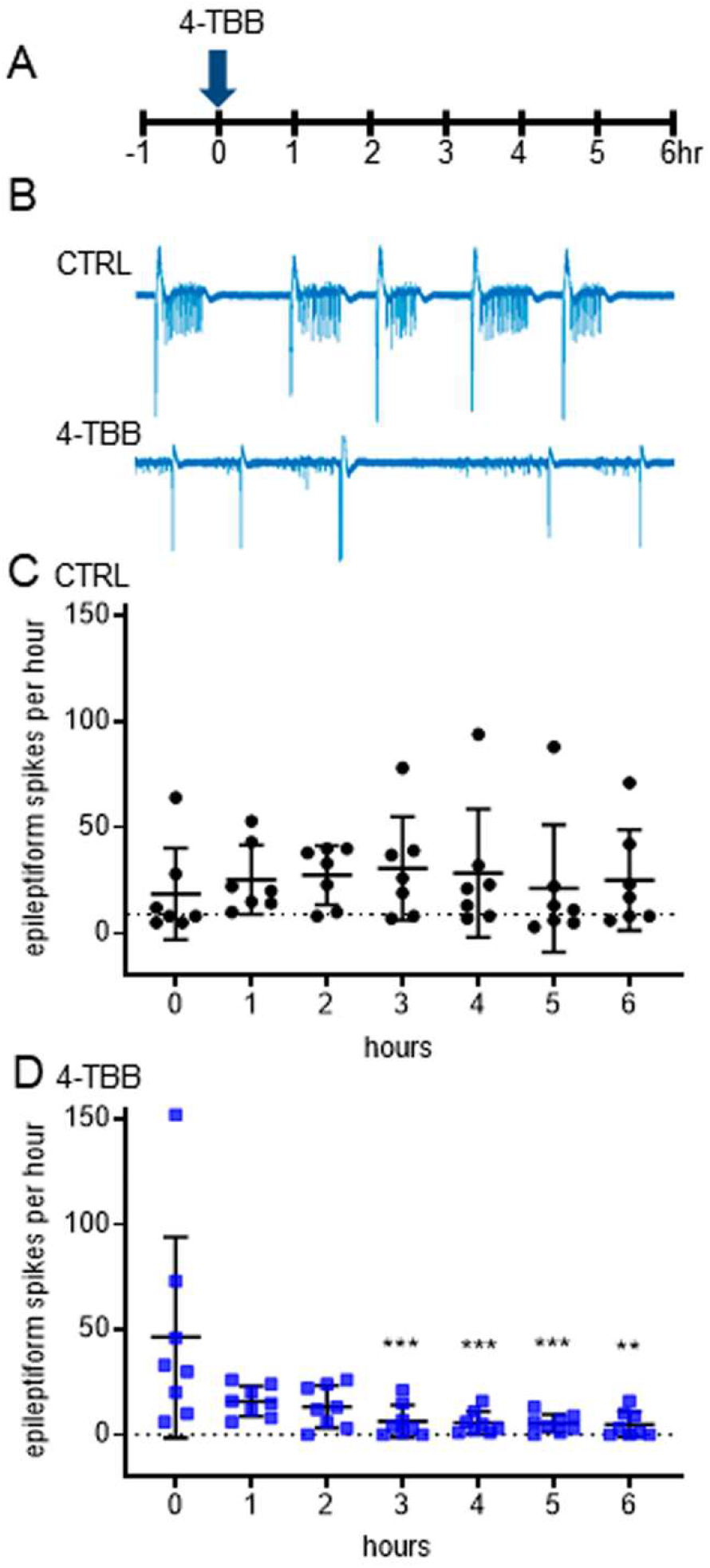
4-TBB is anticonvulsant in mouse brain slice. **A**) Hippocampal slices (7-14 DIV) that exhibited epileptiform activity were exposed to 4-TBB at time ‘0’ during a 7 hr recording period. **B**) Representative extracellular recordings of local field potentials (LFPs) showing seizure activity in un-treated (CTRL) and treated (4-TBB) slices. **C**). Untreated slices (CTRL) show robust epileptiform activity throughout the recording period. **D**) 4-TBB significantly reduces frequency of epileptiform activity, n = 7 (CTRL) and n = 8 (4-TBB) independent slices, respectively. Significance for the effect of 4-TBB *vs*. CTRL was tested using a two-way ANOVA (F_(13, 84)_ = 5.823, *p* < 0.0001). Analysis of the effect of 4-TBB shows that the reduction in epileptiform activity is significant from 3 hrs onwards (F_(13, 91)_ = 2.473, *p* = 0.0063 Šídák’s multiple comparisons test). Bars show means ± SD. ** = *p* < 0 .01, *** = *p* < 0.001.

### 4-TBB reduces induced PTZ-induced seizure in mice

Mice were exposed to 4-TBB (800 mg/kg, s.c. injection) or saline vehicle (CTRL), once per day for 3 days. We observed no overt change in behaviour, nor weight loss, during the test-period. Four hours after the last injection on day 3, seizures were induced by PTZ (60 mg/kg, s.c. injection). Latency to onset of Straub tailing (a first indicator of seizure) was significantly delayed in the 4-TBB-exposed group (215.0 ± 81.0 vs. 138.3 ± 47.3 s, n = 12 and 16, respectively, *p* = 0.004, Fig. 3A). Time to onset of first generalised tonic-clonic seizure was also significantly delayed (239 ± 59 vs. 166.8 ± 44.1 s, 4-TBB vs. saline control, n = 12 and 16, respectively, *p* = 0.001, Fig. 3B). To confirm the expectation that this anticonvulsive effect of 4-TBB was associated with up-regulation of Pum, *post mortem* brains were probed by Western blot. Expression of Pum1 and Pum2 were significantly up-regulated in 4-TBB-exposed mice (1.4 ± 0.3 and 1.2 ± 0.2 fold increase, *p* < 0.0001 and *p =* 0.01, respectively, Fig. 3C) compared to the saline controls (set as 1).

**Figure 3.**
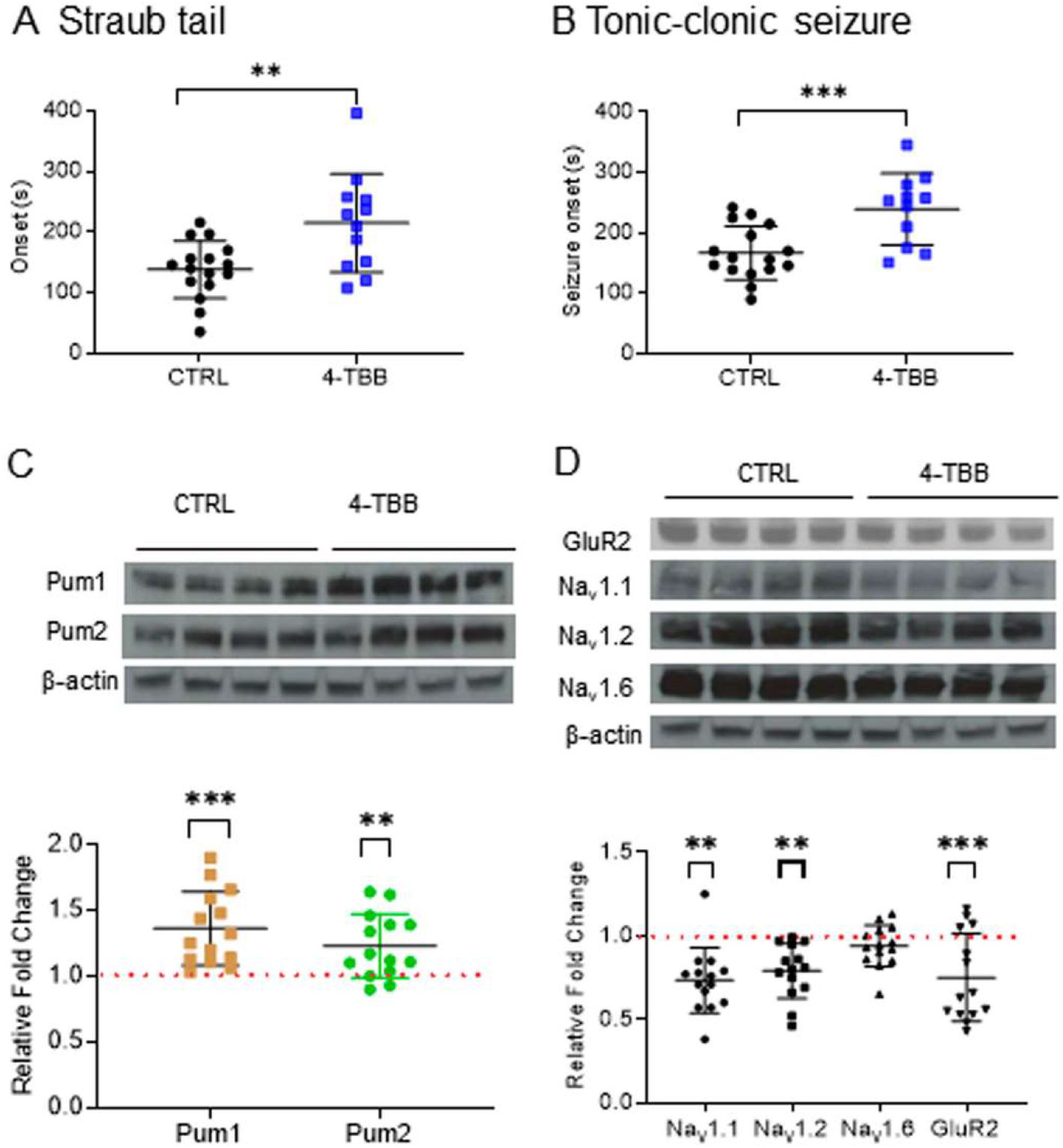
4-TBB is anticonvulsant in the mouse PTZ-induced seizure model. Exposure to 4-TBB increases time to onset of **A**) Straub tail (*p* = 0.004) and **B**) generalised tonic-clonic seizure (*p* = 0.001). **C**). *Post-mortem* analysis of brains, taken from the mice used in the assay, shows expression of Pum1 (*p* < 0.0001) and Pum2 (*p* = 0.01), is up-regulated in animals treated with 4-TBB compared to controls (set as 1). Bars show means ± SD. Inset shows an example Western Blot. Significance was tested in A - C using unpaired *t*-tests. **D**) Western Blot analysis of Na_v_1.1, 1.2, 1.6 and GLUR2 expression in mouse brain show that exposure to 4-TBB is sufficient to down-regulate Na_v_1.1 (*p* = 0.005), 1.2 (*p* = 0.007), and GLUR2 (*p* = 0.001), but not Na_v_1.6 (*p* = 0.75). n = 14, Bars show means ± SD. Significance was tested using ANOVA (F_(4, 65)_ = 6.7, *p* = 0.0001) followed by Dunnett’s post-hoc test (multiple experimental groups). Inset shows an example Western Blot. ** = *p* < 0 .01, *** = *p* < 0.001

Validated Pum-dependent regulated transcripts, in mammals, include *Na*_*v*_*1*.*1 (SCN1A), Na*_*v*_*1*.*6 (SCN8A)* and *GLUR2* (aka *GLUR-A*, AMPA receptor) ^2,8,21^. Bioinformatic analysis of expressed mRNAs also identifies a putative PRE motif in *Na*_*V*_*1*.*2 (SCN2A)*, indicative that this channel variant is also regulated by Pum ^12^. Western blot (Fig. 3D) of the same brain extracts, as above, shows that the expression level of Na_V_1.1, Na_v_1.2 (0.73 ± 0.2 and 0.79 ± 0.16, *p* = 0.005 and 0.007, respectively, n = 14) and GLUR2 (0.75 ± 0.26, *p* = 0.001, n = 14) were significantly reduced in brain tissue exposed to 4-TBB. No change was observed for Na_v_1.6 (0.94 ± 0.12, *p* = 0.75).

### Identification of a more potent 4-TBB analogue

The above data shows proof-of-principle that a compound that effects increased Pum activity has significant potential as an anticonvulsant. However, the active concentration of 4-TBB required for significant effect, in the *in vivo* mouse PTZ-induced seizure assay (800mg/kg) is relatively high compared to other clinically used CNS compounds. Testing 4-TBB at a lower dose (400mg/kg) did not result in statistically significant effects on seizure (data not shown). To identify a more potent analogue, we exploited *Drosophila para*^*bss*^ to screen for anticonvulsive activity of a diverse set of fourteen (synthesised or purchased) 4-TBB analogues (Fig. 4, all structures and chemical properties are shown in sTable 1). We identified 4-(3,5-dimethyl-1*H*-pyrazol-4-yl)benzoic acid (hereafter termed RAB216) to be potently active against *para*^*bss*^ at both 2mM (concentration added to food, Fig. 4A) and at 0.1mM (4-TBB was inactive at this level, data not shown). Analysis of mouse brain exposed to RAB216 (200mg/kg, once per day for 3 days), showed a significant increase in Pum2 expression (*p* = 0.01), and a smaller (but not significant, *p* = 0.07) increase in Pum1 (Fig. 5A). We also screened for effect of valproate (VPA, an effective anticonvulsant in the PTZ-induced seizure assay) and saw no change to either Pum1 or 2 (Fig. 5B). Consistent with its heightened potency to increase Pum2 (*c*.*f*. Fig. 3C), RAB216 was > 4x more active than 4-TBB in protecting mice against PTZ-induced seizures, being significantly active at 200mg/kg (Fig. 5C-D), a dose which prevented the induction of seizure in 50% of animals tested (*p* = 0.05, Fig. 5E). By contrast, a repeat of 4-TBB exposure (800mg/kg), in this assay, prevented tonic-clonic seizures in only 30% of animals during the 20min observation period (not significantly different to CTRL, data not shown). This provides confidence that further, yet more active chemical structures, may be discovered.

**Figure 4.**
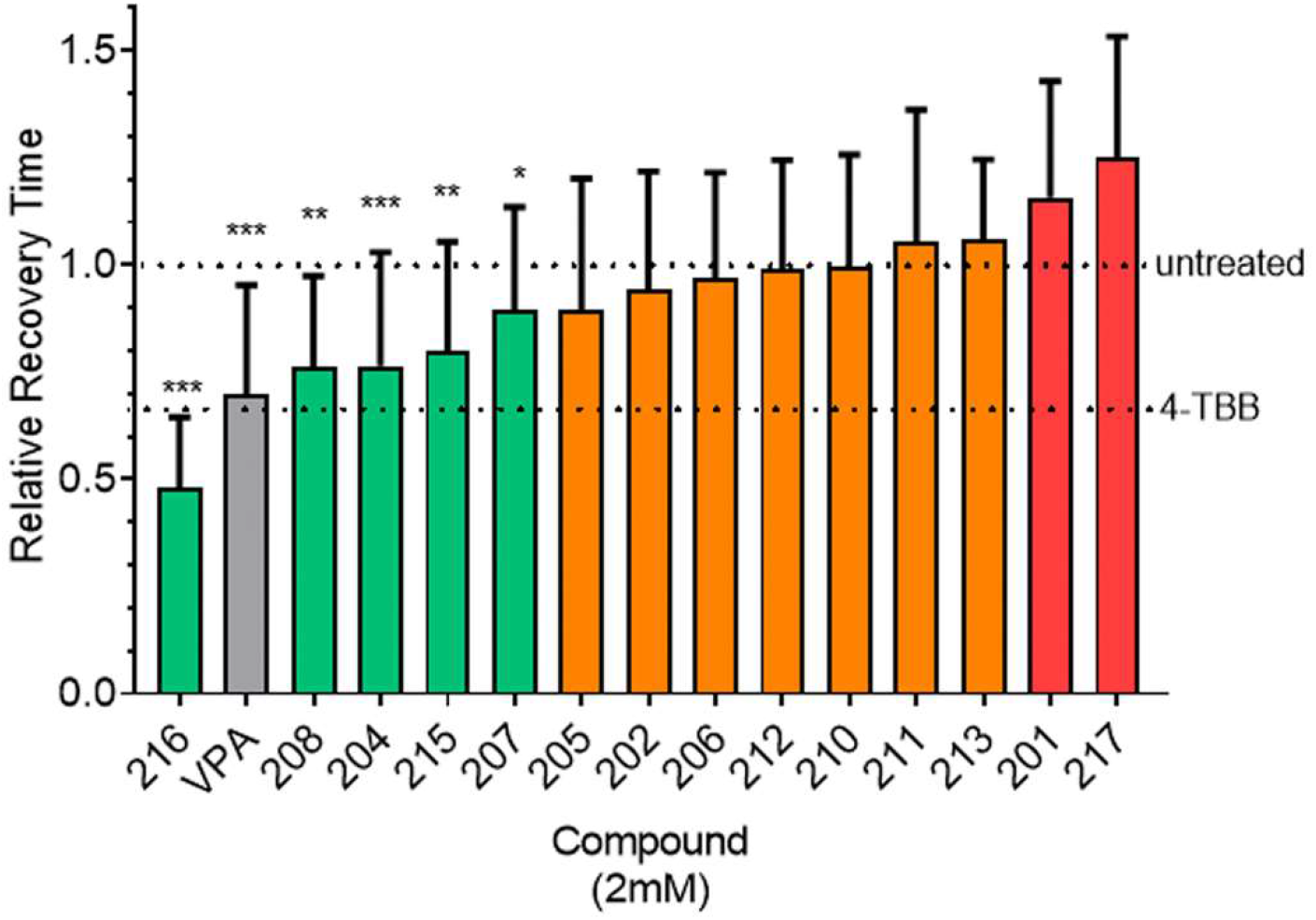
Identification of a more active 4-TBB analogue. 4-TBB analogues (structures shown in sTable 1) identified a number of active compounds, of which 4-(3,5-dimethyl-1*H*-pyrazol-4-yl) benzoic acid (RAB216) was the most potent. Relative recovery time (recovery time normalised to *para*^*bss*^ run at the same time as drugs) was calculated as a ratio of the treatment group (*para*^*bss*^ + compound) recovery time compared to the corresponding untreated group (*para*^*bss*^ – compound) from that week of screening. Green denotes a significant reduction in recovery time, orange denotes no change and red a proconvulsant effect. Sodium valproate (VPA, gray bar) was included as an additional control. * = *p* < 0.05, ** = *p* < 0 .01, *** = *p* < 0.001. (unpaired *t*-tests, between *para*^*bss*^ – compound vs. *para*^*bss*^ + compound).

**Figure 5.**
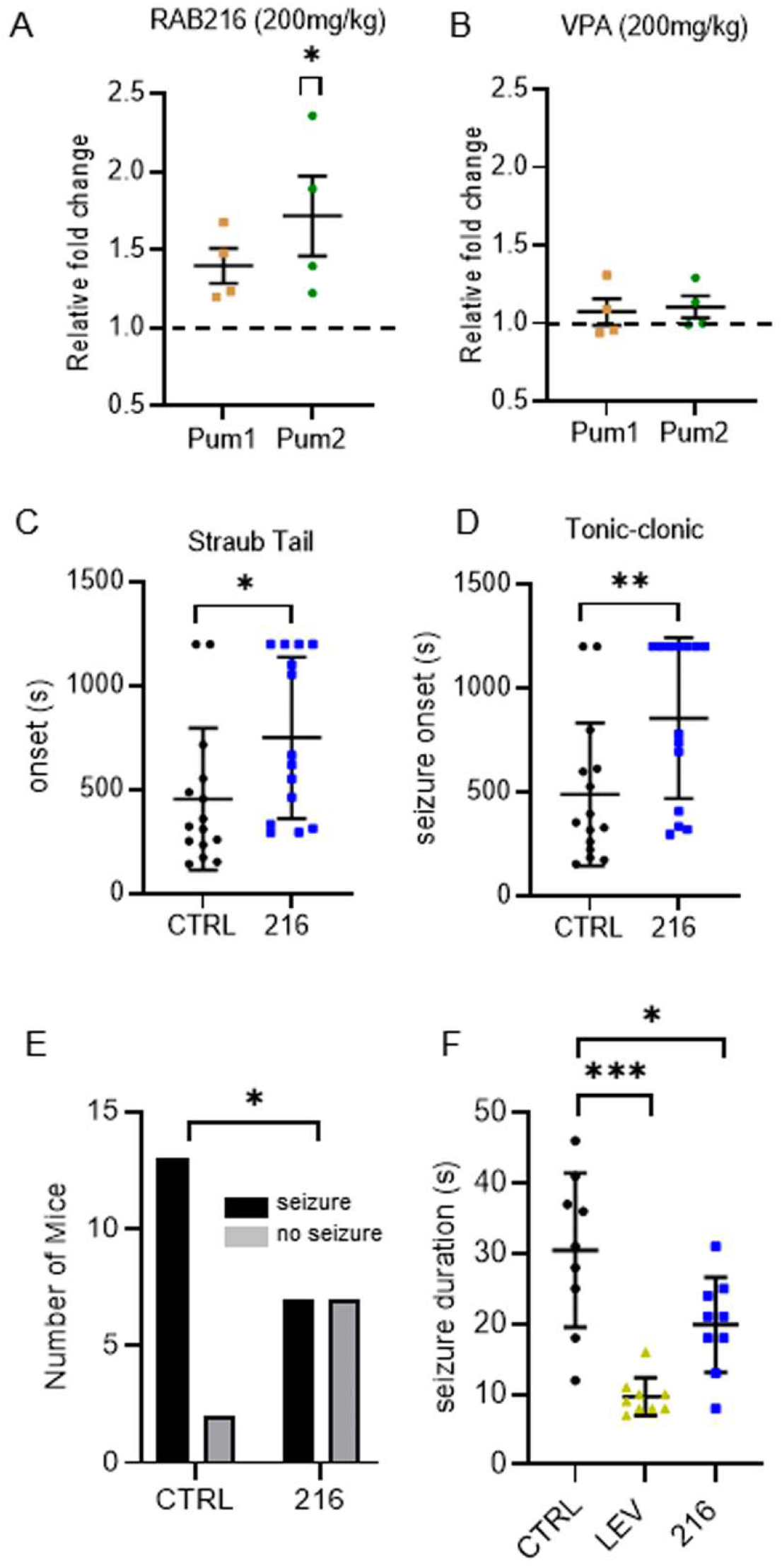
Characterisation of RAB216 shows it to be more potent than 4-TBB **A**) Western blot analysis of the effect of exposure of mouse brain to RAB216 on Pum 1 (n = 4, *p* = 0.07) and 2 (n = 4, *p* = 0.01) expression (unpaired *t*-tests). **B**) Sodium valproate (VPA), which is an effective anticonvulsant in this assay, had no effect on expression of Pum1 or 2 (n = 4, *p* > 0.05). **C**) Effect of RAB216 on Straub tail (*p* = 0.03) and **D**) first appearance of a tonic-clonic seizure (*p* = 0.01) in the PTZ-induced seizure assay. Timings were capped at 20 min (1200 sec). N = 15, (unpaired *t*-tests). **E**) Percentage animals exposed to PTZ showing tonic-clonic seizure following exposure to either saline (CTRL) or RAB216 (*p* = 0.05) respectively, (n = 15 and 14, respectively, Fishers exact test). **F**) Exposure to RAB216 reduces 6Hz electrically-indued seizure duration (n = 9 *p* = 0.013). Levetiracetam (LEV, 100mg/kg, n = 9, *p* <0.0001) was used as a positive control with known efficacy in this assay. Significance was tested using ANOVA (F_(2, 24)_ = 16.88) with Dunnett’s post-hoc multiple comparison. * = *p* < 0.05, ** = *p* < 0 .01, *** = *p* < 0.001.

Finally, we tested RAB216 using a 6Hz electrically-induced seizure assay, which has utility to model drug-refractory epilepsy ^22^. Using the same drug administration protocol as for the PTZ-induced seizure assay, prior exposure to RAB216 (200mg/kg) significantly reduced the length of induced seizure in this assay (*p* = 0.013, n = 9, Fig. 5F). By contrast, 4-TBB (800mg/kg) did not significantly reduce seizure duration (*p* = 0.1, data not shown).

## Discussion

We report here that 4-TBB and analogues, especially RAB216, are potent anticonvulsants with a novel mode of action that involves up-regulation of the homeostatic regulator Pum. Neuronal homeostasis is likely to assume particular importance in epileptic circuits because of the extreme levels of activity associated with the condition. Notably though, homeostasis has never been specifically targeted for anticonvulsant control. An added appeal for targeting homeostatic mechanisms is that these may, realistically, be expected to impact rather less on normal physiology. As such, we predict that anticonvulsant strategies, involving neuronal homeostasis, may prove less susceptible to side effects, which are the primary reason for switching anti-epileptic medication in the clinic ^23^. This is because homeostatic mechanisms have multiple in-built protective regulatory controls that work to prevent under-or over-activation; in the case of Pum, this protein also targets its own mRNA and at least one of its cofactors ^6^.

Whilst multiple forms of neuronal homeostasis have been described, including synaptic scaling and presynaptic regulation of neurotransmitter release ^24,25^, the compounds we describe seemingly manipulate firing rate homeostasis, which acts to maintain action potential firing within pre-determined and physiologically relevant limits ^2^. This is achieved, at least in part, by Pum-dependent control of voltage-gated Na^+^ channel synthesis ^1^. Analyses of Na_v_ protein levels, in *post-mortem* mouse brains, pre-exposed to 4-TBB, validates this mode-of action, showing reduction in Na_v_s 1.1 and 1.2 but, interestingly, no change in Na_v_ 1.6. Intuitively, one might predict a reduction in Na_v_1.6 because gain-of function mutations in the encoding gene, Na_v_ 1.6, are associated with hyperactivity and epilepsy ^26^. By similar logic, the observed reduction of Na_v_1.1 is also unexpected given that this channel type predominates in GABAergic inhibitory neurons ^27^. Our analysis of these known targets of Pum is, however, relatively crude in treating the whole brain as a single tissue. This approach similarly identifies reduced expression of the GLUR2 AMPA receptor subunit following exposure to 4-TBB. Again, how a reduction in this receptor subunit affects neuronal activity, particularly across the entire brain, is difficult to predict. Glutamatergic synaptic currents, in neurons with reduced expression of GLUR2, exhibit increased deactivation rates which may limit the degree of depolarization induced in the postsynaptic cell ^28^. Whilst details remain to be resolved, the changes we observe in Na_v_ and GLUR2 protein levels are consistent with increased Pum expression and, in this regard, serve to strengthen our hypothesis that up-regulation of this homeostatic regulator contributes to the anticonvulsant effect of 4-TBB and RAB216.

We can at present only speculate on how 4-TBB-like molecules mediate an increase in Pum expression. Indeed, in this regard, it is interesting to consider how neurons monitor their activity which, in turn, is transduced to regulate the activity status of intrinsic homeostatic mechanisms. In the case of Pum, we have reported, in *Drosophila*, that synaptic depolarization regulates expression of p300, a histone acetyltransferase that forms a complex with Mef2. As synaptic depolarization increases, levels of p300 reduce, releasing Mef2 from the complex. Once released, Mef2 binds the *dpum* promoter and transactivates gene transcription ^19^. In mammals, by contrast, the level of Mef2 expression is itself activity-regulated, increasing with depolarization ^29^, and analysis of human and mouse *pum2* promoters identifies multiple Mef2 binding motifs ^19^. p300 is also reported to regulate Mef2 in mammals ^30^, but how this protein is influenced by synaptic depolarization has not been described. In mammals, Mef2 also increases the expression of micro-RNAs, including miR-134, which is sufficient to down-regulate expression of Pum2 ^31,32^. Significantly, block of miR-134, using an antagomir, is anticonvulsive in rodents ^33^. Thus, it is conceivable that 4-TBB, and analogues, might act at any level throughout this seemingly complex regulatory mechanism that ensures appropriate expression of Pum proteins. It is expected that levels of Pum are tightly regulated given the requirement to guard against under or over-activity of neuron activity. Indeed, these extensive regulatory and feedback controls, present in Pum-dependent homeostasis, may prove beneficial in exploiting this system for anticonvulsive therapy: minimising potential side-effects of exposure to 4-TBB or its analogues.

In summary, the study we report here provides a first proof-of-principle that manipulation of neuronal homeostasis, and in particular Pum, provides an exploitable route to suppress seizures and, moreover, may be suitable for the treatment of patients that have drug-refractory seizures.

## Acknowledgements

We thank Aoibhinn Kelly, Iona Hayes, Thomas Humphreys and Ceri Hughes who contributed to the 4-TBB structure-activity assay as part of their final year BSc. Projects. This work was supported by funding from the Biotechnology and Biological Sciences Research Council (BB/L027690/1 to RAB); Epilepsy Research UK (PGE2002, Explore Pilot Grant to RAB, RW and SA); an MRC-funded PhD studentship (to FM), a Wellcome Trust-NIH PhD studentship (205944/Z/17/Z to CMGS), the Vrije Universiteit Brussel (to NA and IS) and by MRC (MR/R005427/1 to AT). Work on this project benefited from the Manchester Fly Facility, established through funds from the University and the Wellcome Trust (087742/Z/08/Z).

## Author contributions

Collected data: FM, WHL, CMGS, NA, YNF, GC. RP. Analyzed data: FM, WHL, CMGS, NA, RAB. Supervised experiments: IS, AT, RW, SA, SF, RAB. Wrote manuscript: RB, AT

## Data availability statement

All research data supporting this publication are directly available within this publication.

## Materials and methods

### Animals

*Drosophila* were maintained at 25°C on a 16:8 light/dark cycle. Mice were housed on a 12:12 light/dark cycle at a constant ambient temperature of 21 ± 2°C and given access to water and diet *ad libitum*. All procedures were conducted in accordance with local institutional policies and guidelines. All procedures undertaken at Manchester were approved by the University of Manchester Animal Welfare and Ethics Board and conducted in accordance with project licence authority granted under the Animals (Scientific Procedures) Act 1986.

### Seizure behaviour test in *Drosophila*

Wall-climbing, third-instar larvae (L3), of either sex, were subjected to an electric shock (4V DC, 3s) to induce seizure, with or without previous feeding of compound, as described ^34^. Recovery times (RT) are shown which depict the time taken for larvae to recover, evidenced by a full peristaltic wave and normal locomotion. A cut-off time of 420s was used. For compound-feeding studies, eggs were laid on food containing compound (or vehicle, 0.4% DMSO) and larvae were raised (in the presence of drug) until L3. Where experiments were conducted over a number of weeks (e.g. analogue screen shown in Fig. 4), RT was normalized to the *para*^*bss*^ (without compound) run each week.

### dPum:promoter assay

*A dpum* promoter-GAL4 line ^19^ was crossed to attP24 UAS-luciferase flies^35^. Flies carrying the UAS-luciferase transgene alone were used for background controls. Adult flies were allowed to lay eggs in vials containing food with added compound (or vehicle, DMSO) and to develop to L3. Ten L3 CNSs, of either sex, were placed in 100 μl Promega Glo Lysis buffer for each sample, and 5 independent samples collected. CNSs were homogenized, incubated at room temperature (10 min), centrifuged (5 min), and supernatant transferred to a new tube. 30 μl of each sample was then transferred to a well of a white-walled 96-well plate at room temperature, 30 μl Promega Luciferase reagent was added to each well and plates incubated in the dark (10 min). Luminescence was measured with a GENios plate reader (TECAN, Reading, UK). Values were normalized to total protein concentration, measured using the Bradford protein assay (Bio-Rad, Watford, UK).

### Quantitative RT-PCR

QRT-PCR was performed using a SYBR Green I real-time PCR method (Roche, LightCycler® 480 SYBR Green I Master, Mannheim, Germany) as described ^36^. RNA was extracted from 20 L3 CNSs per replicate, of either sex, using the RNeasy micro kit (QIAGEN, Hilden, Germany). Primer sequences (5’ to 3’) were: *actin-5C* (CG4027), CTTCTACAATGAGCTGCGT and GAGAGCACAGCCTGGAT; *dpum* (CG9755), GCAGCAGGGTGCCGAGAATC and CGCGGCGACCCGTCAACG (forward and reverse, respectively). Relative gene expression was calculated as the 2^−ΔCt^, where ΔCt was determined by subtracting the average *actin-5C* Ct value from that of *dpum*.

### Organotypic slice cultures

Slice cultures were prepared from 5 to 9 day old C57BL/6J mouse pups, of either sex, according to the interface organotypic culture method ^37,38^. Brains were removed and hippocampi dissected and transversely sectioned into 350 μm slices (McIlwain tissue chopper). Slices were plated on polyester membrane inserts (0.4 μm pore) in 6 well culture plates (Corning Costar CLS3450-24EA, Sigma-Aldrich, UK), 2-3 slices per insert, containing 1.2ml of feeding media (50% Minimum Essential Media + GlutaMAX, phenol red and with Earle’s salts (Fisher Scientific, Loughborough, UK), 25 % heat-inactivated horse serum (Sigma, Poole, UK), 21.56 % EBSS, 2 % B27 serum (Fisher Scientific) and 36 mM D-Glucose. Slices were kept at 37°C / 5% CO_2_ and the media replaced the day after plating and then 2-3 times weekly depending on plating density. This method of preparing organotypic hippocampal slice cultures induces spontaneous epileptic-like activity without the need for any pharmacological or electrical provocation. The slicing process mimics a traumatic brain injury which leads to cell death, deafferentation and subsequent axonal sprouting – all of which contribute towards the gradual development of seizure-like activity ^39^.

Local field potentials, at Days In Vitro (DIV) 7-14, were recorded (sampling rate 10 kHz) in CA1 using glass borosilicate patch pipettes (∼1-3MΩ, Harvard Apparatus, Kent, UK) and a Multiclamp 700B (Molecular Devices, CA, USA). Slices were perfused with oxygenated ACSF (125 mM NaCl, 26 mM NaHCO_3_, 10 mM glucose, 3.5 mM KCl, 1.26 mM NaH_2_PO_4_, 2 mM CaCl_2_, 1 mM MgCl_2_) and maintained at 33–36°C for 7 hours. The first hour was used as an activity baseline: slices lacking seizure-like discharges were discarded. After the first hour of baseline activity, slices were bathed in media supplemented with 4-TBB (1.2 mM). All seizure-like events, greater than five seconds, were counted. Analysis was performed using MATLAB (The MathWorks Inc., Natick, MA, USA).

### PTZ seizure-induction

Mice (male, C57BL/6J, 12-15 weeks, 23-30 g,) were injected subcutaneously (sc.) with 0.1 ml of compound (in NaCl, 0.9% w/v saline) or saline vehicle, once per day for 3 days. Four hours after the last injection on day 3, a single dose of PTZ (60 mg/kg/sc. in saline.) was injected. Each mouse was placed into a separate clear plastic arena and videoed for 20 min. After the observation period, mice were anesthetized with isoflurane (3-4% in 20% O_2_ and 50% N_2_O, 0.5 l/min), transcardially perfused with 0.9% saline and brains removed and stored at - 80°C. Seizure scoring was carried out from videos, independently scored by two experimenters blinded to the experimental groups until full analysis was complete.

### 6Hz seizure-induction

NMRI mice (35−45g, male, Charles River, Chatillon-sur-Chalaronne, France) were used. Prior to the electrical stimulation, 0.5% xylocaine was applied to the cornea to induce local anesthesia and ensure good conductivity. Corneal stimulation (46 mA, 0.2 ms duration pulses at 6 Hz for 3 s) was administered by a constant current device (ECT Unit 57800; Ugo Basile, Comerio, Italy) ^40,41^. Acutely evoked 6-Hz seizures were characterized by stun, forelimb clonus, twitching of vibrissae, and/or Straub-tail. For each animal, the total seizure duration was manually recorded. Ip administration of levetiracetam (LEV, 100mg/kg) 1h before seizure induction was used as a positive control ^42^. Seizure scoring was carried out live during the experiment. The entire experiment was also video recorded. The researcher was blinded to the experimental groups until full analysis was complete.

### Western blot

Whole brain was homogenised in ice-cold buffer (150 mM NaCl, 50 mM Tris-HCl, 1% Nonidet P-40, 0.5% Sodium deoxycholate and 0.1% SDS) containing protease inhibitors (Promega, Madison, USA) and centrifuged at 10,000× g (30 min at 4°C). Supernatant was stored at −20°C. Antibodies were: anti-Pum 1 (1:1000, #12322, Cell Signaling, MA, USA), anti-Pum 2 (1:1000, ab10361, Abcam, Cambridge, UK), anti-SCN1A (1:1000, ASC-001, alomone labs, Jerusalem, Israel), anti-SCN2A (1:1000, ASC-002, alomone labs), anti-SCN8A (1: 1000, ASC-009, alomone labs), anti-GluR2 (1:2000, AB1768-I, Merck, Darmstadt, Germany) and anti-β-Actin (1:5000, ab8227, Abcam). Samples (25 μg of protein) were separated by SDS page, and protein transferred to a polyvinylidene difluoride membrane (GE Healthcare). After blocking (0.5 % BSA and 0.05% TWEEN-20 in Tris-buffered saline, TBS-T), membrane was incubated overnight (4°C) in primary antibody diluted in 0.5% BSA in TBS-T. Membranes were incubated with HRP-conjugated secondary antibodies (1:2500, #7074, Cell Signaling) in 0.3% BSA in TBS-T and blots developed with an Enhanced Chemiluminescent Detection Kit (Pierce, Rockford, USA). Protein band density was measured using Image J (NIH, USA).

### Chemical synthesis

RAB216 was designed by first screening analogues of 4-TBB and its carboxylic acid derivative RAB102 (data not shown). The structure activity relationship (SAR) revealed that only compounds with a carboxylic acid or aldehyde group directly bonded to the benzene ring were active, even when replaced with isosteric groups. Furthermore, a change of the benzene ring to a pyridine or indole ring, or a change of substituents from para to meta, resulted in complete loss of activity. Electronegative groups were tolerated in the para-position, increasing the likelihood of a compound forming strong intermolecular bonds with its binding partner. Therefore, when designing the second-generation compounds (sTable 1), some analogues included electronegative oxygen and nitrogen atoms. This included RAB216 which contained a pyrazole ring, providing extra interactions to enhance activity. Additionally, analysis of the screen showed that compounds with an element of 3D structure were more active: in RAB216 the two methyl groups attached to the pyrazole group caused it to be twisted by approximately 18°. Reviews of drug libraries suggest that planar compounds are less likely to be biologically active ^43^. Compounds, including RAB216, were designed in accordance with Lipinski’s rules, such as molecular weight (under 500) and lipophilicity (logP under 5), to give desirable physiochemical properties ^44^. All of the compounds screened, including RAB216, fit within these guidelines (sTable 1).

### Statistics

Statistical significance was tested using either a Student’s two-tailed *t*-test (paired or unpaired), a one-way or two-way ANOVA followed by post-hoc testing (multiple experimental groups) or Fishers exact test. Level of significance on figures is indicated by * (*p* ≤ 0.05), ** (*p* ≤ 0.01), *** (*p* ≤ 0.001). Figures show means ± SD.

