## Supplemental Table 1 for "Targeting neuronal homeostasis to prevent seizures"

| Compound | Structure | MW (g/mol) | CLogP | Polar surface area Å² | pKa | Synthesised / Purchased (supplier + cat no.) |
|---|---|---|---|---|---|---|
| **4-TBB** | 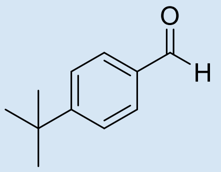 | 162.23 | 3.49 | 17.07 | N/A | FluoroChem: 065159 |
| **RAB201** | 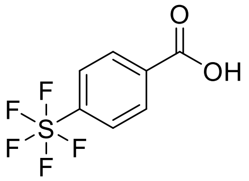 | 247.99 | 3.12 | 37.30 | 3.8 | Merck: ATC311764220 |
| **RAB202** | 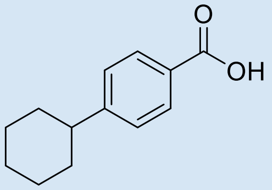 | 204.11 | 4.51 | 37.30 | 4.3 | Merck: AOBH961DC0B4 |
| **RAB204** | 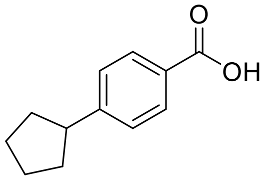 | 190.09 | 3.95 | 37.30 | 4.3 | Merck: AOBH97EBAF06 |
| **RAB205** | 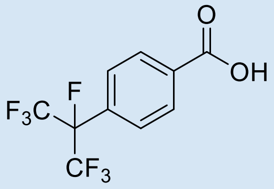 | 290.01 | 3.56 | 37.30 | 3.0 | Merck: ENA514506754 |
| **RAB206** | 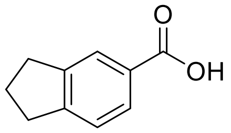 | 162.06 | 2.90 | 37.30 | 4.1 | Fluorochem: 059793 |
| **RAB207** | 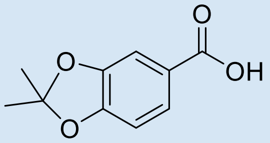 | 194.05 | 2.61 | 55.76 | 3.9 | Merck: COMH04239142 |
| **RAB208** | 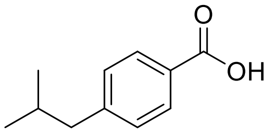 | 178.09 | -3.84 | 37.30 | 4.2 | Fluorochem: 238558 |

| | | | | | | |
|---|---|---|---|---|---|---|
| **RAB210** | 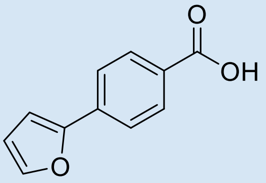 | 188.04 | 3.16 | 46.53 | 4.3 | Fluorochem: 031925 |
| **RAB211** | 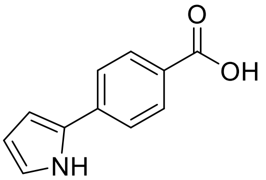 | 187.06 | 2.64 | 49.33 | 4.4 | Synthesised |
| **RAB212** | 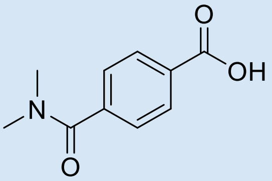 | 193.07 | 0.61 | 57.61 | 4.2 | Synthesised |
| **RAB213** | 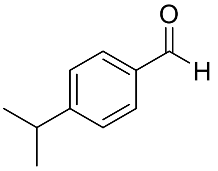 | 148.08 | 2.92 | 17.07 | None | Fluorochem: 225406 |
| **RAB215** | 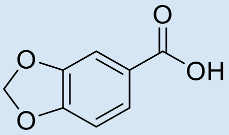 | 166.02 | 1.97 | 55.76 | 3.8 | Fluorochem: 022643 |
| **RAB216** | 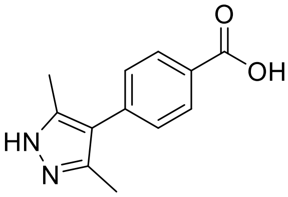 | 216.24 | 1.92 | 61.7 | 4.0 | Synthesised |
| **RAB217** | 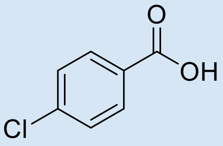 | 156.57 | 2.70 | 37.3 | 4.0 | Fluorochem: 118580 |

sTable 1: Structure and physiochemical properties of compounds screened and source of compounds (supplier or synthesised in house).
